# All–atom molecular dynamics simulation of the combined effects of different phospholipids and cholesterol contents on electroporation

**DOI:** 10.1101/2022.07.21.501058

**Authors:** Fei Guo, Ji Wang, Jiong Zhou, Kun Qian, Hongchun Qu, Ping Liu, Shidong Zhai

## Abstract

The electroporation mechanism could be related to the composition of the plasma membrane, and the combined effect of different phospholipids molecules and cholesterol contents on electroporation is rarely studied and concluded. In this paper, we applied all-atom molecular dynamics (MD) simulation to study the effects of phospholipids and cholesterol contents on bilayer membrane electroporation. The palmitoyl-oleoyl-phosphatidylcholine (POPC) model, palmitoyl-oleoyl-phosphatidylethanolamine (POPE) model and 1:1 mixed model of POPC and POPE called PEPC were three basic models. An electric field of 0.45 V/nm was applied to nine models including three basic models with cholesterol contents of 0%, 24%, and 40%. The interfacial water molecules moved under the electric field, and once the first water bridge formed, the rest of the water molecules would dramatically flood into the membrane. The simulation showed that a rapid rise in the Z component of the average dipole moment of interfacial water (Z-DM) indicated the occurrence of electroporation, and the same increment of Z-DM represented the similar change in the size of water bridge. With the same cholesterol content, the formation of the first water bridge was the most rapid in POPC model regarding the average electroporation time (*tep*), and the average *tep* of the PEPC model was close to that of the POPE model. We speculate that the difference in membrane thickness and initial hydrogen bonds of interfacial water affecting the average *tep* among different membrane composition. Our results reveal the influence of membrane composition on electroporation mechanism at the molecular level.

## 1 Introduction

When cells are exposed to an electric field, pores form in the cell membrane and increase its permeability, which is known as electroporation. ^1–3^ The technologies based on electroporation were applied in medicine, biotechnology and even food processing, ^4–8^ but the electroporation mechanisms have not been well understood. Various theoretical models have been developed to investigate electroporation at the level of single cell, multiple cells, and tissues. ^9–15^ However, traditional electroporation simulations were difficult to observe the dynamic process of electroporation from a microscopic perspective.

All-atom molecular dynamics (MD) simulation is an efficient tool which can directly observe the occurrence of electroporation at the molecular level. ^16,17^ Tieleman found that the water molecules moved with the electric field gradient and continuously entered the cell membrane when a vertical electric field was applied. ^18^ Hu et al. constructed a dipalmitoyl-phosphatidyl-choline (DPPC) bilayer model and found that the anode side of the phospholipid head first shifted and formed voids under the electric field, and water molecules entered the membrane through the voids to form water bridges. ^19^ However, Vernier and Ziegler found that the deflection of the headgroup dipole would not directly lead to the formation of water bridges at the nanoscale, whereas the field-oriented rotation of the headgroup dipole, the water dipole, and the solvation interaction were important reasons for the formation of electroporation. ^20,21^ Additionally, researchers found that the rearrangement of water molecules at the water / membrane boundary of a POPC phospholipid bilayer, the magnitude and direction of the interfacial water dipole moment were the keys to the formation of electroporation, which further revealed the electroporation mechanisms. ^22–25^ Furthermore, some researchers found that with the movement of water molecules, the hydrogen bonds between water would change, and the network of hydrogen bonds vibrated, which affected the occurrence of electroporation. ^26–28^ The variation in hydrogen bonds between water affected the recombination of peptides on the membrane, and the association between protein peptides and cell membranes had been linked to diseases such as Alzheimer disease. ^29–31^

Most all-atom MD models used to study electroporation mechanism were based on pure phospholipid membrane models. ^32–34^ However, the presence of different phospholipid molecules in phospholipid membrane could change its stability and fluidity, which in turn affected the electroporation. ^35–37^ Different phospholipid molecules influenced the shape of the phospholipid membrane and thus regulated protein properties. ^38^ Different lipid enzymes were enriched in different phospholipid molecules, which changed the physicochemical properties of the membrane. ^39^ These factors made the difference of different phospholipid membranes under electric field to some extent. Gurtovenko applied the same electric field to the POPC phospholipid bilayer and the POPE phospholipid bilayer, respectively, and found that the POPE model required more time for the occurrence of electroporation, compared to the POPC model. ^40^ Polak et al. found that the architecture of lipid head groups and tails affected the poration propensity in membranes containing fluid phase lipids. ^41^ Furthermore, they compared archaeal lipids (glucosyl inositol and inositol headgroup) with normal DPPC lipids and found that the electroporation threshold of DPPC was larger at the same temperature. ^42^ Hu et al. found that the architecture of lipid head groups and tails in coarse-grained martini membranes were also the vital reason determined the poration propensity, similarly as in atomistic membranes, and the higher poration propensity was in fully saturated lipids rather than monounsaturated lipids. ^43^

Cholesterol is also an important component of phospholipid membrane and the influence of cholesterol content on electroporation of simple lipid bilayers had been proposed. ^44–46^ In recent years, the researchers found that the presence of cholesterol could significantly change the thickness, ^47–52^ and the fluidity of plasma membrane which in turn affected the breakdown voltage of the membrane. ^47,53–55^ Cholesterol increased lipid cohesion and hardness to prevent peptide-induced damage, reducing the rate of membrane damage, but the effect was even reversed for different phospholipids. ^56,57^ In experiments, cholesterol was commonly used for vesicles. The researchers used cholesterol to prepare giant unilamellar vesicles, and found that the rupture of the plasma membrane under the electric field would be hindered by cholesterol. ^37,58–60^ Most simulations suggested that cholesterol delayed the occurrence of electroporation, but the effect of cholesterol on the perforation propensity of lipid bilayer was strongly dependent on the structure of lipids. Mauroy et al. reported that the decrease of poration propensity in POPC phospholipid was due to the increase content of cholesterol, ^61^ but Portet found that the increasing cholesterol accelerated the occurrence of electroporation in dioleoylphosphocholine (DOPC) vesicles. ^62^ Recently, Kramar added the 20%, 30%, 50% and 80% content of cholesterol on the POPC phospholipid, and found that when the content was too high, the breakdown voltage of the membrane would not increase but decrease. ^63^ Ruiz-Fernández et al. found that under nanosecond pulsed electric field (nsPEF), the phospholipid membrane with high cholesterol content would have significant conformational changes in the voltage sensing domain (VSD) of *Ga*^2+^ channel, which affected the movement of water molecules. ^64^ In conclusion, the influence of cholesterol and different phospholipids on electroporation mechanism remains to be studied.

The combination of phospholipid molecules and cholesterol enriched plasma membrane diversity. However, most electroporation simulations and experiments with cholesterol did not consider its interaction with different phospholipid molecules. The POPC phospholipid molecules existed in the cell membranes of brain cells, erythrocyte, which were one of the important phospholipids in biophysical experiments. ^65,66^ The POPE phospholipid molecules were also widely found in the cell membranes of erythrocyte and in Escherichia coli. ^67^ As two main phospholipid components in mitochondrial membrane, the POPE phospholipids and the POPC phospholipids were often used in studies under electric field respectively. ^68,69^ The two phospholipids had similar structure and same tail base, and high sensitivity to electric fields, which facilitated us to observe the dynamic process of electroporation. Therefore, we constructed POPE, POPC, and the 1:1 mixed model of the two (PEPC) as three basic models, and then added 24% and 40% contents of cholesterol. A constant electric field with a magnitude of 0.45 V/nm in the Z-direction was applied in all models. We showed the dynamic process of the formation of water bridge, proposed the relationship between the Z-DM and electroporation. The average time of the occurrence of electroporation (*t*_*ep*_) of each model was calculated, and it was considered that the membrane thickness and the initial hydrogen bonds of interfacial water were the main factors affecting the average *t*_*ep*_. The hydrogen bonds between interfacial water molecules and phospholipid molecules in the mixed models was also calculated. The entire study provides an efficient basis for future electroporation simulations and experiments.

## 2 Models and methods

### 2.1 Phospholipid membrane models

This study included 9 phospholipid membrane models, all of which were constructed by charmm-gui. ^70,71^ The water / membrane / water phospholipid bilayer was composed of phospholipid molecules and TIP3P water molecules. The TIP3P water molecule was simple, stable and widely used in the study of electroporation. More importantly, the TIP3P model had an electric charge and the dipole moment can be calculated quickly, which is suitable for our study. ^33,72^ The phospholipid membrane model composed of single-component of POPC, the phospholipid membrane model composed of single-component of POPE, and the 1:1 mixed model of POPC and POPE (PEPC) were the three basic models for our all-atom MD simulations. The positions of POPC molecules and POPE molecules were randomly distributed in the upper and lower layers of the phospholipid model. Cholesterol molecules were added to the basic models with 24% and 40% contents. The total number of phospholipid molecules and cholesterol molecules in all models was 125, and the total number of water molecules was always 8875. The specific parameters such as the number of molecules of the models were shown in Table. 1. All the models were placed in a cuboid box with the same length in the X and Y directions. The initial box size of the POPE model was 6.3 nm × 6.3 nm × 11.2 nm, and in order to better achieve the balance, the initial box will be slightly changed according to the structure of different models.

**Table 1.** Simulation models for this study

| Model | $N_{POPC}$ | $N_{POPE}$ | $N_{CHL}$ | CHL content (%) | $N_{Water}$ |
| --- | --- | --- | --- | --- | --- |
| 1 | 125 | 0 | 0 | 0 | 8875 |
| 2 | 0 | 125 | 0 | 0 | 8875 |
| 3 | 63 | 62 | 0 | 0 | 8875 |
| 4 | 95 | 0 | 30 | 24 | 8875 |
| 5 | 0 | 95 | 30 | 24 | 8875 |
| 6 | 48 | 47 | 30 | 24 | 8875 |
| 7 | 75 | 0 | 50 | 40 | 8875 |
| 8 | 0 | 75 | 50 | 40 | 8875 |
| 9 | 38 | 37 | 50 | 40 | 8875 |

$N_{POPC}$ = the number of POPC phospholipid molecules
$N_{POPE}$ = the number of POPE phospholipid molecules
$N_{CHL}$ = the number of cholesterol molecules
$N_{Water}$ = the number of water molecules
Model 3, model 6 and model 9 are the mixed model called PEPC

### 2.2 MD simulation

All-atom MD simulations were performed on the NAMD software, and the VMD software was used to observe the dynamic process of molecular simulations and calculate relevant parameters. ^73^ We referred to the protocol of equilibrium for membranes from charmm-gui. ^74^ Using 3D periodic boundary conditions, the temperature was set to 310 K. The whole motion formula was integrated by Velocity Verlet integral method. Long-range electrostatic forces were calculated by Particle Mesh Ewald (PME). The space cut-off distance of the van der Waals force and the longrange electrostatic force was set to 1.2 nm, and the switching function was adopted to slowly reduce to zero. ^33^ The charmm36 force field was applied to the simulations of lipid bilayers, the conjugate gradient method was applied to search for energy successively from the initial gradient until energy minimization. The SHAKE algorithm was used to constrain the bond length of hydrogen atoms and made water molecules rigid. In order to better balanced the entire phospholipid bilayer, the Z direction position of the heavy atoms and the distance of dihedral angles were constrained in a short time. ^75^ Langevin dynamic temperature control method and the Langevin Piston Nose-Hoover pressure control method maintain the temperature and pressure at 310 K and 1 atm, respectively. ^76^

The whole simulation process had the following stages: First, after the minimization of energy, a heating process of 250 ps was performed to increase the temperature to 310 K with a time step of 1 fs. Then, the system was kept at constant temperature and pressure for 1.625 ns under the NPT ensemble, and the time step was changed to 2 fs. Afterwards, the position restriction of the heavy atom of the phospholipid molecule and the restriction of the dihedral angle distance of the phospholipid molecule were removed, and a 10 ns equilibrium process was performed. Finally, a constant electric field of 0.45 V/nm was applied to the models after 10 ns equilibrium. As the water bridge became too large, the model became extremely unstable. ^77^ Therefore, the end time of each model simulation was different. The required time of the whole simulation process is shown in Fig. 1.

**Fig. 1.**
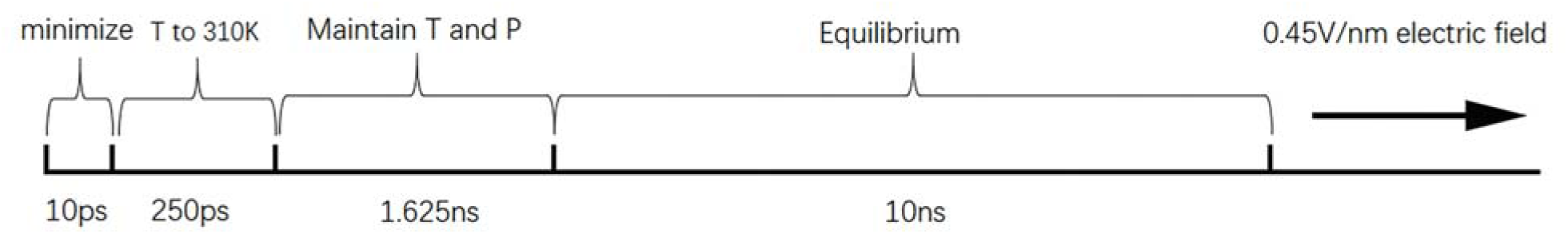
Time spent in each stage of the simulation, T stands for temperature, and P stands for pressure.

## 3 Results and discussion

### 3.1 The dynamic process of electroporation

We first calculated the potential energy and the area per lipid (APL) of PEPC model during the 10ns equilibrium process. Fig. 2 showed that the potential energy curve was relatively stable throughout the 10 ns equilibrium periods. The maximum and minimum value of APL were 0.635 *nm*^2^ and 0.58 *nm*^2^, respectively, and the curve of APL was stable, which were similar to the previous literature studies. ^75,78,79^ The APL and the potential energy in other models were also stable like Fig. 2 during the 10 ns equilibrium.

**Fig. 2.**
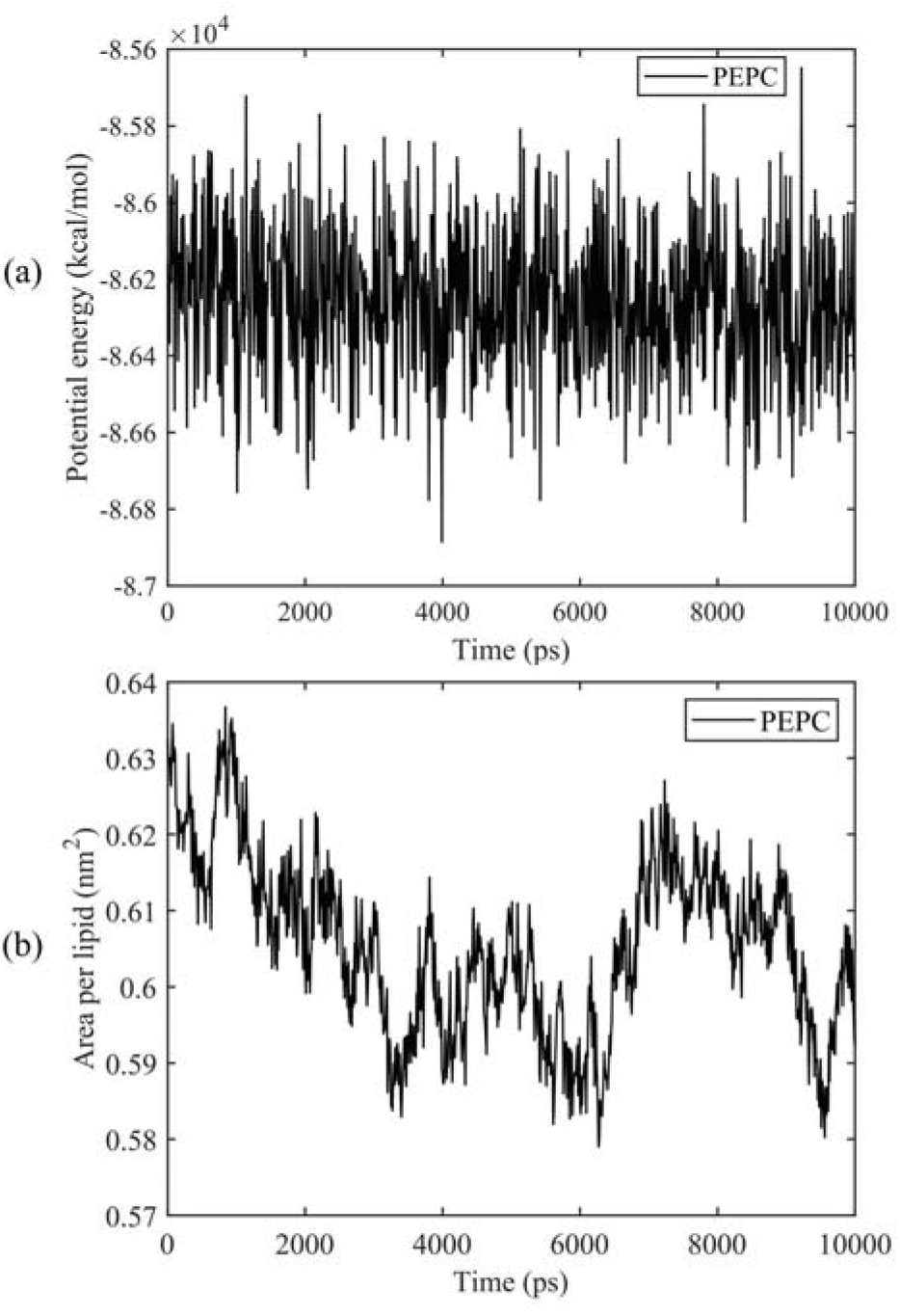
The potential energy and the area per lipid (APL) of PEPC model during the 10 ns equilibrium. (a) Potential energy. (b) APL.

Through all-atom MD simulations, we can observe the dynamic process of electroporation from a microscopic point of view. The process by which water molecules on both sides of the membrane enter the membrane and join to form a water bridge is defined as electroporation. ^33^ Fig. 3 showed the formation of water bridge in the PEPC model. After the 10 ns equilibrium process, the water / membrane / water interface reached a relative stable state, all water molecules were evenly distributed on the outer side of the phospholipid bilayer. The head of the phospholipid molecules moved with the electric field, providing enough space for the water molecule to move, creating a water protrusion. Then, the water molecules continued to move until the water on the upper and lower sides were connected in the interior of membrane, and the water protrusion changed into a water bridge in this process. It is worth noting that during the process in the formation of water bridge, phospholipid molecules will also bend to the water bridge. When the first water bridge formed, its width increased rapidly, multiple water bridges appeared one after another, and water molecules polarized rapidly. The dynamic process of electroporation was similar to most studies. ^32,80,81^ The tendency of nanopore formation process in all other models were similar to that in Fig. 3, but various in times required for water protrusion and bridge formation among different models.

**Fig. 3.**
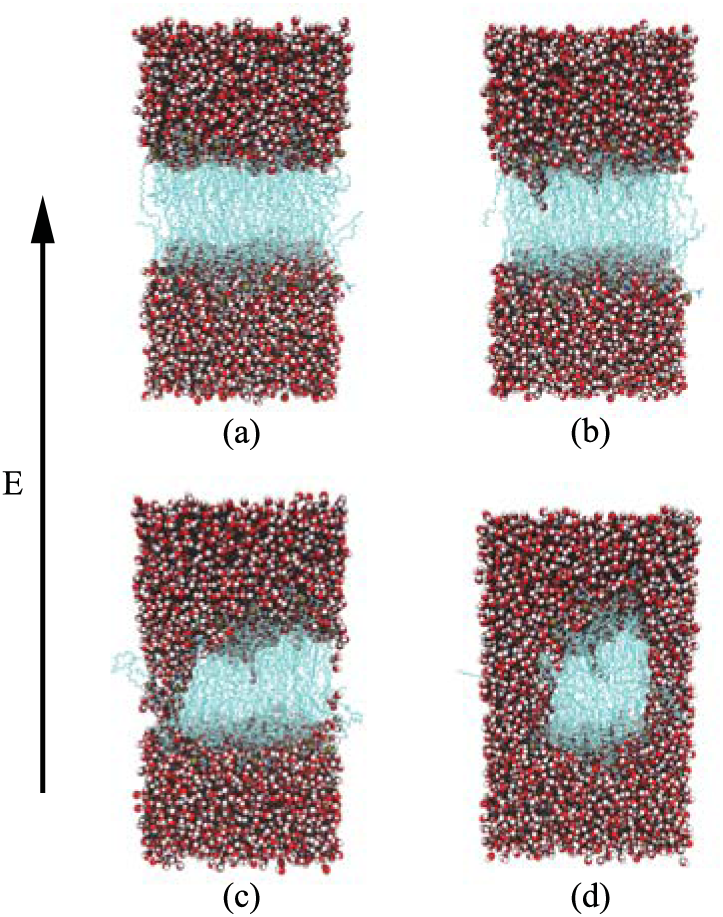
Dynamic process of electroporation in PEPC model under the electric field. The oxygen and hydrogen atoms were represented by red and white spheres, the gold spheres represented the phosphorus atoms of the phospholipid head group, and the blue strings thread in the middle were the phospholipid molecules. (a) 0 ns – initial stage. (b) 0.58 ns – water protrusion. (c) 1.15 ns – water bridge. (d) 1.35 ns – after water bridge.

### 3.2 The relationship between dipole moment and electroporation

The water molecules within 4 Å of the membrane / water interface were defined as the interfacial water, and the movement of interfacial water molecule was an important reason for the formation of water bridge. ^22^ Dipole moment was the most intuitive parameter to reflect the polarization degree of water molecules. ^24^ When interfacial water moved under the electric field and formed water bridges, the hydrogen bonds between water molecules would also change. ^27^ We calculated the Z component of average dipole moment of the interfacial water (Z-DM) and hydrogen bonds between the interfacial water (H-bonds) under electric field. The Z-DM was given by the Z component of total dipole moment, then divided by the number of interfacial water molecules, and the cut-off angle and the cut-off distance of H-bonds were 35° and 3.5 Å. Fig. 4 clearly showed that both the Z-DM and the H-bonds increased exponentially, and the models would become extremely unstable when the water bridges were too large, so the end time of each simulation was different. ^77^ From Fig. 4, the POPC, PEPC, POPE models formed the first water bridge in turn, and the time of PEPC model formed the first water bridge was close to that of POPE. The primary amines of intra and intermolecular hydrogen bonds in POPE head group had a lower poration propensity compared with that in POPC head group. As a result, the POPE model was more intensive which delayed the formation of the first water bridge. ^40^ It is worth noting that the rapid rise in the Z-DM represented the formation of the first water bridge which was also called the occurrence of electroporation.

**Fig. 4.**
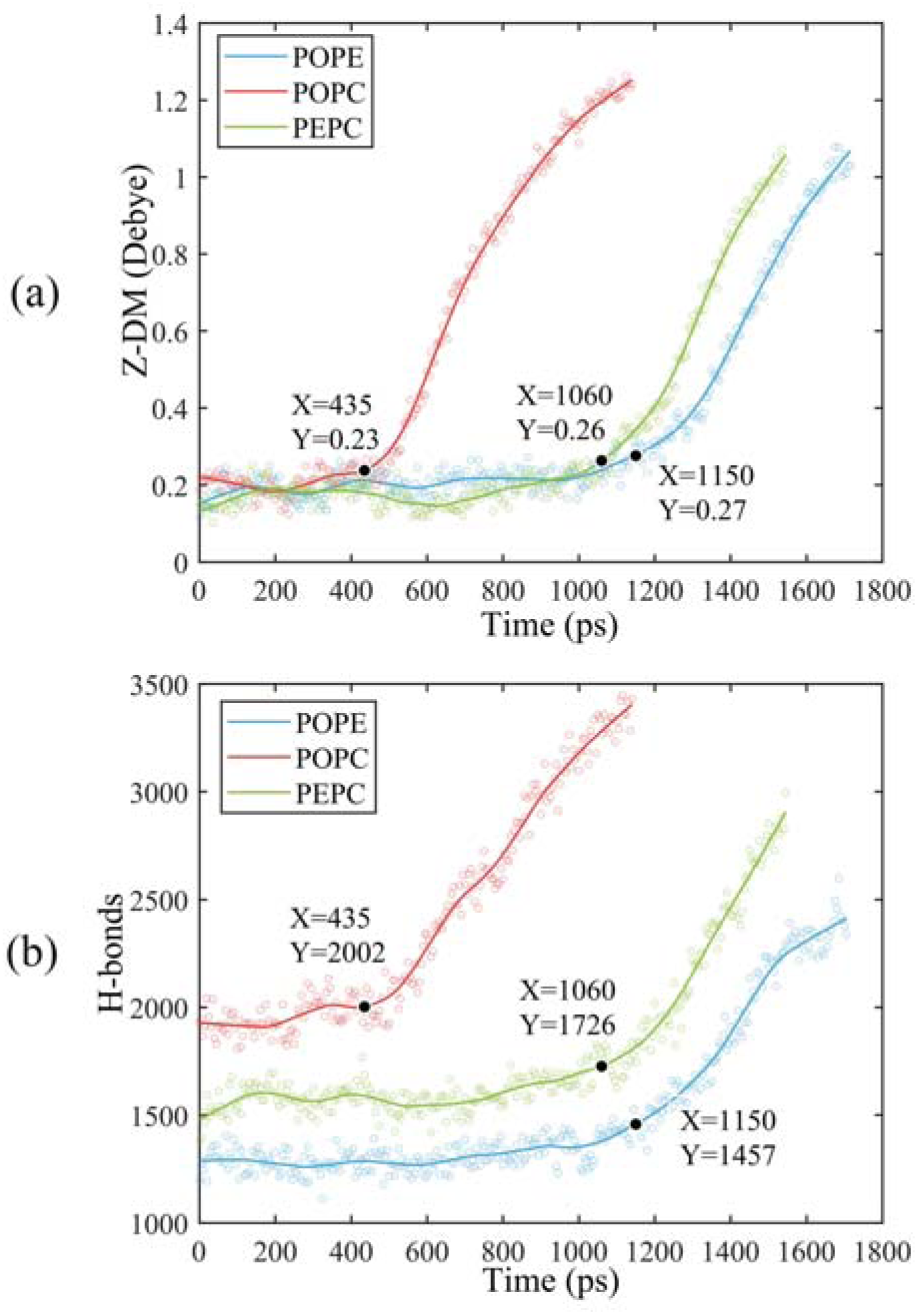
The Z-DM and the H-bonds in POPE, POPC and PEPC model. The scatter plots were our data. In order to facilitate observation, we fitted the data to get the curves, and marked the time when the first water bridge of POPC, POPE and PEPC model formed. (a) Z-DM. (b) H-bonds.

Once the first water bridge formed, the water molecules rapidly flooded into the membrane, which was the reason for the rapid rise of the Z-DM. When the first water bridge of the three models formed, they had similar dipole moment values, 0.23 Debye (D), 0.26 D and 0.27 D. We speculate that the Z-DM is an important parameter to measure whether the first water bridge is formed or not. More kinds of phospholipid molecules and components will be presented in future work to enrich the complexity of models and to approach the real cell membrane, and we consider that Z-DM will also be an indicator to measure the formation of the first water bridge. In Fig. 4(b), the H-bonds also increased rapidly after a period of time. At the moment of water bridge formation, the water molecules were in close contact and the H-bonds rose. The H-bonds of the POPC rose the fastest, PEPC was the second, and POPE was the slowest. Different from Z-DM, the initial number of H-bonds were quite different, there were about 1900 in POPC model, 1500 in PEPC model, and 1300 in POPE model, and the difference was due membrane lipid composition. Therefore, we speculate that the initial number of H-bonds number is also a reason affecting electroporation mechanism. H-bonds in water is an important factor.

From Fig. 5, we recorded the moments corresponding to the same Z-DM among POPE, POPC and PEPC models, and the curves of which were approximately linear and similar in slop. The top view was used to observe the degree of electroporation, which was the size of the water bridge. When the Z-DM was 0.4 D, the corresponding moments in the POPE and POPC models were 1.32 ns and 0.56 ns, respectively, tiny water bridge appeared in both models. When the Z-DM was 0.6 D, which corresponded to the moments of 1.42 ns and 0.65 ns in the POPE and POPC models, respectively, similar obvious water bridge outline was observed in both models. When the dipole moment was 0.8 D, the corresponding moments of the POPE and POPC models were 1.52 ns and 0.73 ns, the water bridge continued to grow, and the similar large pores were generated in both models. In addition, we also gave a top view of the PEPC model. Although the water bridge formed at the edge of the model, it could still be found that the water bridge increased with the Z-DM. According to the above results, the models had similar sizes of water bridge when the magnitude of Z-DM were the same.

**Fig. 5.**
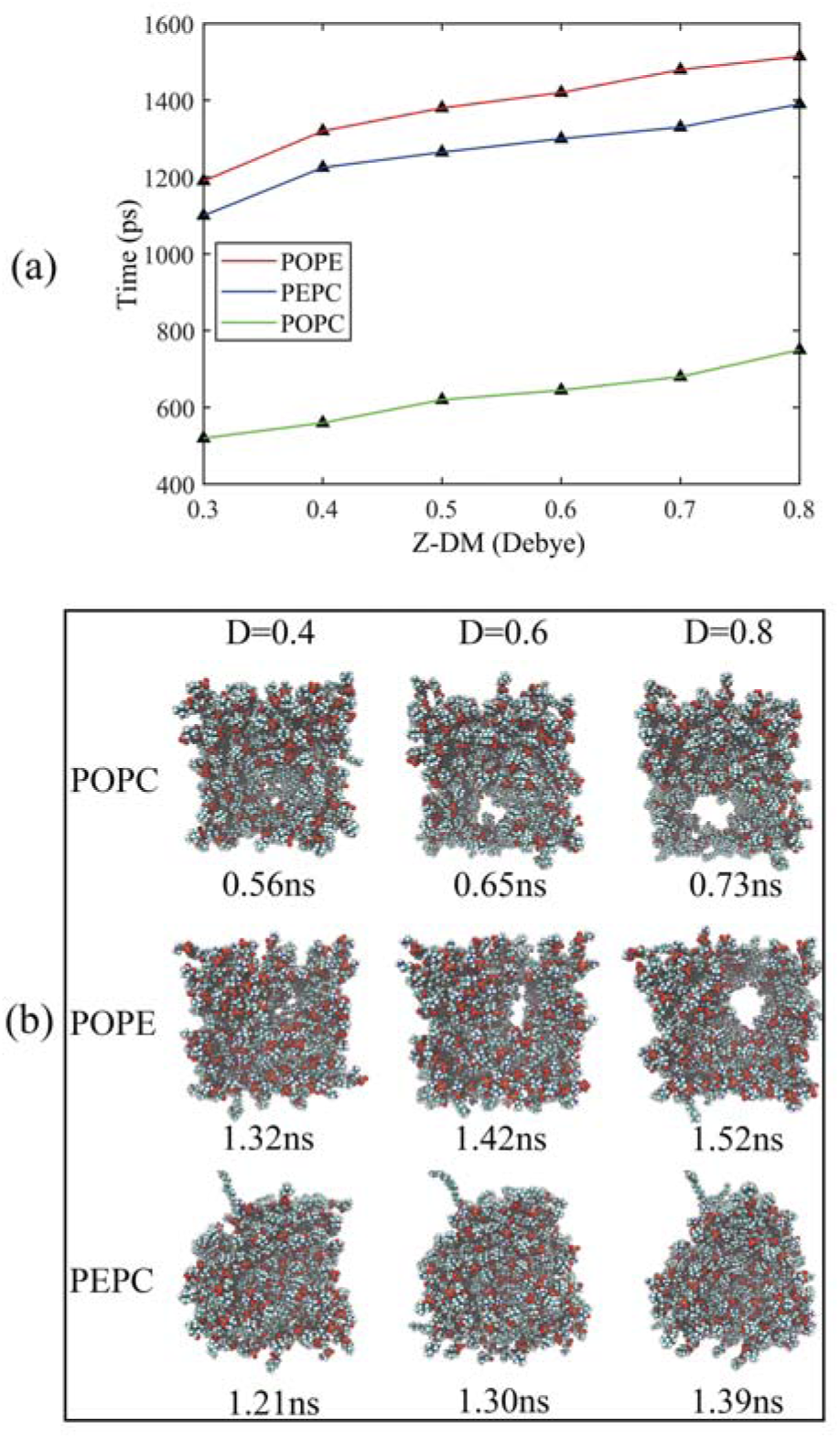
The moments and their corresponding top views of the three models under the same Z-MD. In order to observe the size of water bridge, the top views only showed the phospholipid molecules and hid the water molecules. (a) The corresponding time of the three models at same Z-DM. (b) Top view of the three models at same Z-DM.

We also analyzed the Z-DM for the models containing cholesterol, as shown in Fig. 6. The Z-DM increased exponentially after the occurrence of electroporation, which was similar to the trend in Fig. 4. At the same cholesterol content, POPC, PEPC and POPE model formed water bridge successively. The increase of cholesterol content delayed the formation of water bridge among the same phospholipid models, which was consistent with the simulation result made by Casciola et al. ^82^ In models with cholesterol, the moments of rapid rise in Z-DM were close to the moments of the formation of water bridges, which was consistent with the result in Fig. 4. We recorded the moment when each model had the same Z-DM magnitude of 0.6 D, and gave its corresponding top views. However, when the Z-DM magnitude was the same, the sizes of water bridges were not similar. Unlike the cholesterol-free phospholipid models, the same Z-DM magnitude hardly reflected the same degree of electroporation. In order to found the relationship between the Z-DM and electroporation, we showed the moment and the top views of the Z-DM increment of 0.4 D after the occurrence of electroporation. At the same Z-DM increment of 0.4 D, the sizes of water bridges of 6 models were relatively similar. Then, we also calculated the same Z-DM increments of 0.5 D and 0.6 D, and the results were the same as for the increment of 0.4 D, which further reflected the relationship between the Z-DM increment and the degree of electroporation.

**Fig. 6.**
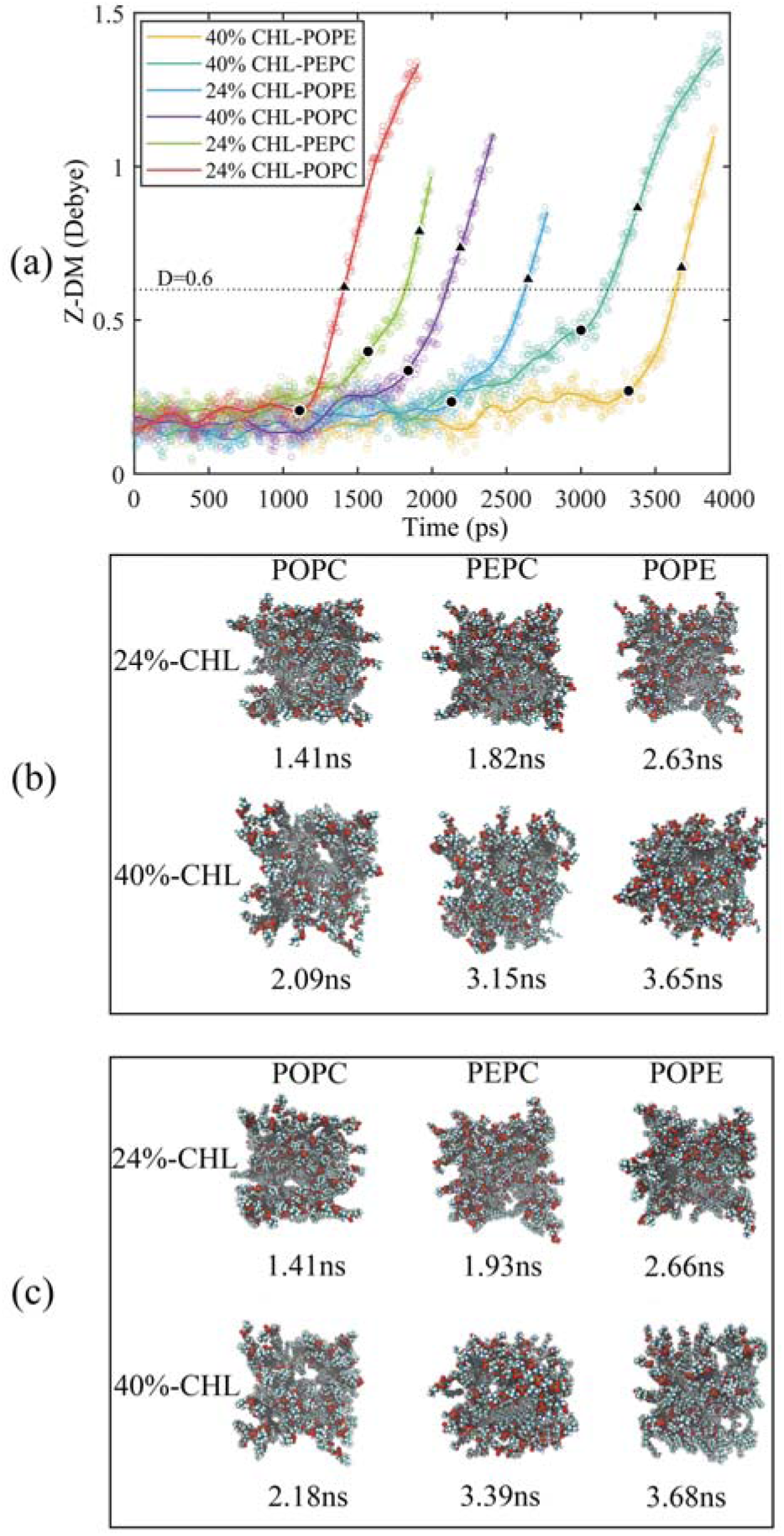
The Z-DM and the top view in cholesterol-containing models. (a) The changes of Z-DM with time. The scatter plots were our data, we fitted the data to get the curves, and the solid circle marked the time when the first water bridge of POPC, POPE and PEPC model formed. The dotted line indicated that the Z-DM was 0.6 D. From the occurrence of electroporation, the increment in Z-DM of 0.4 D were marked by triangle. (b) When the Z-DM was 0.6 D, the top views in all models and their corresponding moments. (c) The moments and corresponding top views of all models when the Z-DM increment was 0.4 D.

### 3.3 Effects of initial hydrogen bonds and membrane thickness on electroporation

The average *t*_*ep*_ were the average values of 10 repeated results for each model, and the membrane thickness of all models were calculated before the electric field. From Fig. 7, with the same cholesterol content, the average *t*_*ep*_ of POPE models were always the biggest, followed by PEPC models, and POPC models were the smallest. The average *t*_*ep*_ of PEPC models was consistently close to that of POPE models rather than POPC models at the three cholesterol contents, but the difference between them rose with the increase in cholesterol. At 0% cholesterol, the average *t*_*ep*_ of PEPC models was 0.1 ns faster than that of POPE models; at 24% cholesterol, the average *t*_*ep*_ of PEPC models was 0.23 ns faster than that of POPE models; and at 40% cholesterol, the average *t*_*ep*_ of PEPC models was 0.77 ns faster than that of POPE models. With the increase in cholesterol content, the average *t*_*ep*_ of our models increased, and this trend of change was similar to the previous study. ^82^ The average *t*_*ep*_ among three basic models of POPC, PEPC and POPE at 0% cholesterol were 0.45 ns, 1.15 ns and 1.25 ns, the average *t*_*ep*_ at 24% cholesterol were 1.11 ns, 1.57 ns and 1.87 ns, and the average *t*_*ep*_ at 40% cholesterol were 1.84 ns, 3.00 ns and 3.32 ns, respectively. The combination of cholesterol molecules and phospholipid molecules will limit the thermal movement of phospholipid molecules to a certain extent, thereby affecting their deflection under the electric field. The similar phenomenon was also shown in Fig. 7(b). At the same cholesterol content, the POPE phospholipid model had the largest membrane thickness, the POPC membrane thickness was the smallest, and the PEPC membrane thickness was among the two. The membrane thickness also increased with cholesterol. The higher cholesterol content is, the less the influence on membrane thickness. The influence of cholesterol on the thickness of membrane was negatively correlated with its concentration. Membrane thickness had been studied for its effect on electroporation mechanism, as with APL, from a structural perspective of the models. ^50,52^ At the same electric field, the thinner phospholipid membranes were more likely to form water bridge, ^43^ and models with larger membrane thickness required larger transmembrane voltages, resulting in delay the average *t*_*ep*_. At the same time, a small membrane thickness corresponds to a large APL, ^80^ which means that the space between the phospholipid molecules widen, and it is easier for water molecules to enter the membrane and electroporation. In our model, the difference in membrane thickness was not particularly large, but it also clearly leaded to the difference in *t*_*ep*_. In the next step in our study, we will compare phospholipid combinations with larger membrane thickness differences, such as POPC and DMPC, where the membrane thickness of DMPC is much less than that of POPC. This will better confirm our conclusions.

**Fig. 7.**
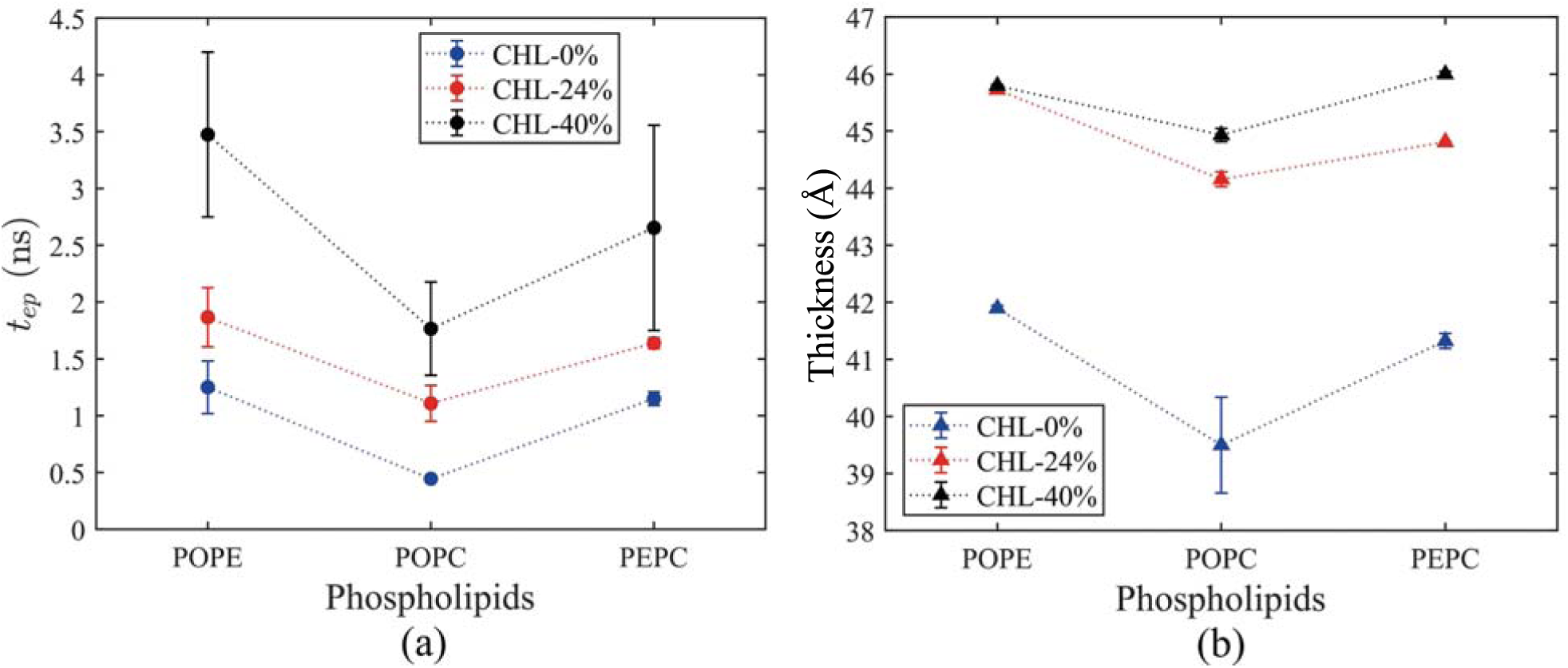
Average *t*_*ep*_ and membrane thickness of all models. The average *t*_*ep*_ and membrane thickness were the average values of 5 repeated simulation results for each model, the membrane thickness was calculated before the electric field. The dotted lines were made to observe the changing trend. (a) Variation of average *t*_*ep*_ over time. (b) Variation of membrane thickness over time.

Fig. 8(a) showed the initial H-bonds of all models. The initial H-bonds was the average of interfacial water H-bonds at 10 ns equilibrium, which means that the initial H-bonds was only relevant to the composition of model. With the increase in cholesterol content, the initial H-bonds of the three basic phospholipid models decreased. At the same cholesterol content, the initial H-bonds of POPC, PEPC and POPE model decreased successively. According to Table. 2, when the cholesterol content ranged from 0% to 24%, the change rates of average *t*_*ep*_, membrane thickness and initial H-bonds of interfacial water were 146%, 12% and -25%, respectively. When cholesterol content ranged from 24% to 40%, the change rates of average *t*_*ep*_, membrane thickness, and initial H-bonds of interfacial water were 67%, 2%, and -14%, respectively. From the rate of change, it seems that the initial H-bond is also a factor that affects average *t*_*ep*_, we speculated that both the membrane thickness and the initial H-bonds affected the average *t*_*ep*_. From Fig. 8(b), within the first 1 ns of applying the electric field, the average H-bonds of the POPC, PEPC, and POPE models with 40% cholesterol content were 1243, 1097, and 939, and the less the H-bonds in the first 1 ns, the later the electroporation occurred. The H-bonds were relatively stable at first, but when the electroporation occurred, it suddenly rose exponentially. The rapid rise in H-bonds also indicated the formation of water bridge, which is a same conclusion in Fig. 4(b). At 24% cholesterol content, we can also draw a similar conclusion of that in 40% cholesterol content. H-bonds has been applied to describe the process of the models from equilibrium state to the formation of water bridges, ^27,28^ but the initial H-bonds was not considered. The combination of other phospholipid molecules (POPS and POPG) with cholesterol also had some influence on the H-bonds, ^83^ which verified our results of different initial H-bonds. The part of the simulation results are useful for our next stage of work. The combination of different phospholipid molecules and cholesterol content in phospholipid membrane changes the thickness and initial H-bonds all of which influenced the average *t*_*ep*_.

**Fig. 8.**
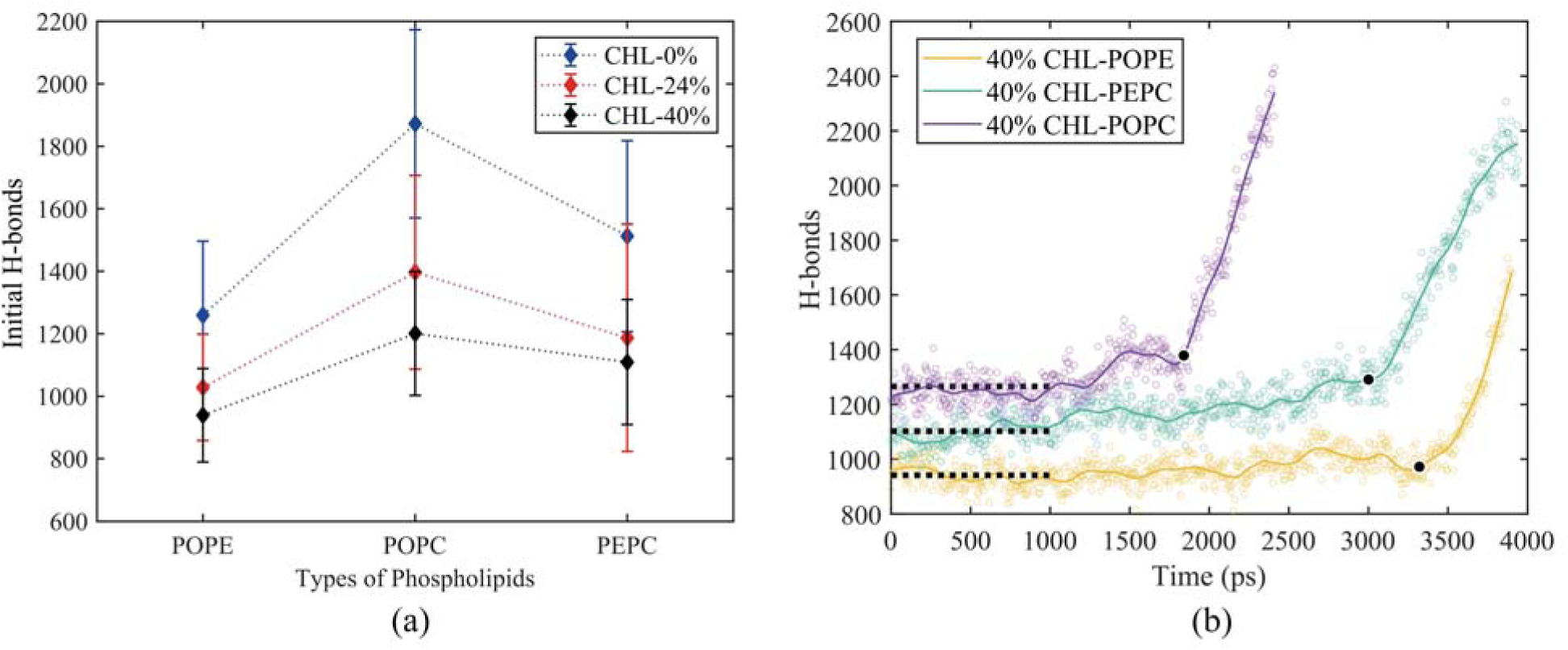
The initial H-bonds and the H-bonds of 40% cholesterol content models under the electric field. The initial H-bonds was the average value of the H-bond at the 10 ns equilibrium. (a) Initial H-bonds of all models. (b) The H-bonds of the 40% cholesterol models under the electric field. The black dots represent the moment when electroporation occurs. The dotted lines represent the average of interfacial water H-bonds over the first 1ns electric field applied.

**Table 2.** The average *t*_*ep*_, membrane thickness and initial H-bonds in POPC models.

| CHL-content (%) | Time (ns) | Membrane thickness (Å) | H-bonds |
| --- | --- | --- | --- |
| 0 | 0.45 | 39.49 | 1872 |
| 24 | 1.11 | 44.15 | 1396 |
| 40 | 1.77 | 44.93 | 1200 |

To further understand the influence of different phospholipid molecules on electroporation, we calculated the H-bonds between phospholipid molecules and interfacial water in the PEPC model. As shown in Fig. 9, the H-bonds between phospholipid molecules and interfacial water decreased with the increase of cholesterol content. At the contents of 0%, 24%, and 40% of cholesterol, the H-bonds between POPC phospholipid molecules and the interfacial water (H-bonds of POPC-water) were always greater than those between POPE phospholipid molecules and interfacial water (H-bonds of POPE-water). At 0% cholesterol, the difference between the number of H-bonds of POPC-water and that of POPE-water was 103; at 24% cholesterol, the difference between the number of H-bonds of POPC-water and that of POPE-water was 97; at 40% cholesterol, the difference between the number of H-bonds of POPC-water and that of POPE-water was 87. Under the electric field, water molecules are close to the POPC molecules rather than the POPE molecules. At the same time, we speculated that this was the reason why the average *t*_*ep*_ of POPC model was shorter than that of POPE model.

**Fig. 9.**
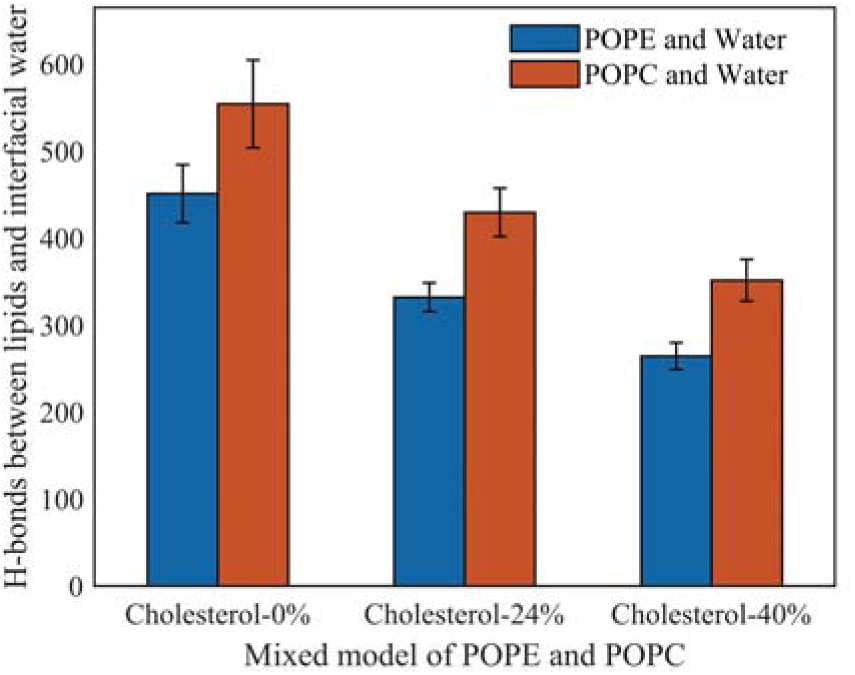
Hydrogen bonds between phospholipid molecules and interfacial water in PEPC model under the electric field at three cholesterol concentrations.

## 4 Conclusions

In this study, we used all-atom MD simulations and systematically investigated the influence of different phospholipids and cholesterol content on electroporation. Under the electric field, water molecules entered the membrane through the gaps and formed water protrusion. The water protrusion would become water bridge after a period of time. Interestingly, the occurrence of electroporation was accompanied by the rapid rise of the Z-DM and the H-bonds. From the top view of the phospholipid models, we found that the magnitude of Z-DM could reflect the size of water bridge, which was also called the degree of electroporation. However, the water bridges of different models were no longer similar with the same Z-DM after adding the cholesterol, while the sizes of water bridges in different models had the similar size with the same increment in Z-DM.

With the same cholesterol content, POPC models had the shortest average *t*_*ep*_, POPE models had the longest average *t*_*ep*_, and the average *t*_*ep*_ of PEPC models were close to those of POPE models. The overall average *t*_*ep*_ decreased with the increase of cholesterol content, but the order of average *t*_*ep*_ remained unchanged for the three basic phospholipid models with the same cholesterol content. We calculated the parameters that vary with the model composition, the initial H-bonds and the membrane thickness. At the same cholesterol content, the POPC model had the smallest membrane thickness, the POPE model had the largest membrane thickness, membrane thickness of PEPC model was between the two. The initial H-bonds of POPC, PEPC and POPE model decreased successively. The presence of cholesterol reduced the initial H-bonds and increased the membrane thickness. By combining the variation trend of initial H-bonds and membrane thickness with that of average *t*_*ep*_, we inferred that membrane thickness and initial H-bonds were important factors affecting average *t*_*ep*_. The influence of different phospholipid membranes and cholesterol content on electroporation mechanism were studied from the molecular level. The effect of dipole moment on the formation of water bridge and the effect of initial H-bonds and membrane thickness on average *t*_*ep*_ were proposed. In conclusion, this paper provides a reference for the future all-atom MD research of electroporation mechanism.

## Acknowledgements

This work was supported in part by the Science and Technology Research Program of Chongqing Municipal Education Commission (Grant No.KJQN202100607), in part by the Natural Science Foundation of Chongqing, China (cstc2020jcyj-msxmX0393), and in part by the National Natural Science Foundation of China (No. 51507024).

